# Kappa Opioid Receptors in Mesolimbic Terminals Mediate Escalation of Cocaine Consumption

**DOI:** 10.1101/2023.12.21.572842

**Authors:** L. Gordon-Fennell, R.D. Farero, L.M. Burgeno, N.L. Murray, A.D. Abraham, M.E. Soden, G.D. Stuber, C. Chavkin, L.S. Zweifel, P.E.M. Phillips

## Abstract

Increases in drug consumption over time, also known as escalation, is a key behavioral component of substance use disorder (SUD) that is related to potential harm to users, such as overdose. Studying escalation also allows researchers to investigate the transition from casual drug use to more SUD-like drug use. Understanding the neurobiological systems that drive this transition will inform therapeutic treatments in the aim to prevent increases in drug use and the development of SUD. The kappa opioid receptor (KOR) system is typically known for its role in negative affect, which is commonly found in SUD as well. Furthermore, the KOR system has also been implicated in drug use and importantly, modulating the negative effects of drug use. However, the specific neuronal subpopulation expressing KOR involved has not been identified. Here, we first demonstrated that pharmacologically inhibiting KOR in the nucleus accumbens core (NAcC), as a whole, blocks cocaine escalation under long-access self-administration conditions. We then demonstrated that KOR expressed on ventral tegmental area (VTA) neurons but not NAcC neurons is sufficient for blocking cocaine escalation by utilizing a novel virally-mediated CRISPR-SaCas9 knock-out of the *oprk1* gene. Together, this suggests that activation of KOR on VTA terminals in the NAcC drives the transition to the SUD-like phenotype of escalation of cocaine consumption.

## INTRODUCTION

Substance use disorder (SUD) and harmful drug consumption is an ongoing epidemic, with currently no treatment options for cocaine use disorder specifically. One key component of SUD is the transition from casual to compulsive drug-taking, often coupled with an increase in overall drug consumption which leads to a higher likelihood of harm to the user. This SUD-like phenotype, termed escalation, can be modelled in animals utilizing a long-access (six-hour) drug self-administration (SA) paradigm, allowing researchers to probe the neurobiological processes underlying this transition to inform future therapeutics^1^.

The kappa opioid receptor (KOR) / dynorphin system is implicated in components of SUD including stress^2–4^, withdrawal^5,6^, and overall negative affect^7^. Cocaine itself increases prodynorphin mRNA in the nucleus accumbens (NAc)^8,9^, a brain region implicated in SUD that receives dense dopaminergic innervation from the ventral tegmental area (VTA). Dopamine dynamics, particularly in the NAc, are associated with the rewarding and reinforcing properties of most addictive substances^10–12^. Activation of KOR, a Gi-coupled receptor, inhibits dopamine release in the NAc^13^. Both medium spiny neurons in the NAc and VTA dopamine terminals express KOR^14–17^. The ability of KOR to modulate dopamine release in the NAc and KOR’s role in aversive phenotypes underlines its potential to be a key system in the transition to compulsive drug-taking. Prior studies have already demonstrated that pharmacological inhibition of KOR systemically and locally in the NAc shell blocks escalation of heroin^18^ and methamphetamine^19^ consumption. However, the subpopulations of KOR within the NAc responsible for this effect have yet to be studied.

In this paper, we examined the role of KOR in the NAc core (NAcC) in the progression of cocaine escalation. We utilized the long-lasting KOR antagonist, norBNI, to pharmacologically inhibit KOR in the NAcC during long-access cocaine SA. To determine if KOR mediates effects through pre-synaptic or post-synaptic activation, we also utilized a novel CRISPR-SaCas9 construct targeting the *oprk1* gene to knock-out KOR from NAcC medium spiny neurons or VTA neurons prior to long-access cocaine SA. Through these experiments, we found that KOR activation on VTA terminals in the NAcC, but not medium spiny neurons of the NAcC, is necessary for escalation of cocaine consumption.

## METHODS

### Subjects

Male and female Wistar rats were purchased from Charles River and singly housed on a 12-hr light-dark cycle (lights on at 7:00) with chew-toy enrichment and *ad libitum* access to laboratory chow and water. Upon arrival, male rats weighed 250-275g and female rats weighed 200-225g. All behavioral experiments were performed during the light cycle. The pharmacological inhibition of KOR with norBNI was performed solely in male rats. All other experiments were performed in both female and male rats. All animal use and procedures were approved by the University of Washington Institutional Animal Care and Use Committee.

### Surgical Procedures

Animals were acclimated in the housing facility for at least one week before surgical procedures were performed. For rats that underwent multiple surgeries, there was a minimum of one week recovery period between the surgeries. All surgeries were performed utilizing aseptic technique. Anesthesia was induced and maintained with isoflurane (1-5%). An NSAID (Meloxicam, 1mg/ml/kg, in 3mL total saline, s.c.) was administered on the day of surgery and post-operative day 1. Surgical areas were shaved and cleaned with alternating ethanol and betadine.

#### Cannula Implantation

Custom 26G bilateral guide cannulas with a 2.6mm center-to-center distance and 7mm projection were purchased from Plastics One (8IC235G26XXC). The rat was fixed in a stereotaxic frame (Kopf), the head was leveled, and burr holes were drilled overlaying our target region. We lowered the guide cannula 1mm dorsal to the target region (**NAcC** angle: 0°, AP: 1.3mm, ML: +/- 1.3mm, DV: -7.2mm) and secured the implant to the skull with dental cement. A dummy cannula with 0mm projection (Plastics One) was inserted and a dustcap (Plastics One) was screwed onto the guide cannula to block contaminates.

#### Viral Injection

The rat was fixed in a stereotaxic frame (Kopf), the head was leveled, and burr holes were drilled overlaying our target regions. A Hamilton syringe (7105KH 5.0ul SYR (24/2.75”/3), ref#: 88000) operated by Micro4 Microsyringe Pump was used to deliver the viral vector (1μL per site at 250nL/min). We injected a 1:8 viral cocktail of AAV1-Cre-GFP and AAV1-FLEx-SaCas9-U6-HA-sgOprk1 (*oprk1* knock-out CRISPR) or AAV1-FLEx-SaCas9-U6-HA-sgGabrg2 (control CRISPR). We lowered the Hamilton syringe 0.2mm ventral to the target region (**NAcC** angle: 0°, AP: 1.3mm, ML: +/- 1.3mm, DV: -7.2mm ; **VTA** angle: 0°, AP: -6.35mm, ML: +/- 0.5mm, DV - 8.5mm; distances measured from bregma), waited for 2min, began the 4min injection while raising the syringe to 0.2mm dorsal to the target DV over the first minute, and then waited 8min post-injection before slowly raising the syringe out of the brain. At the conclusion of the surgery, incisions were closed with sutures and cleaned with betadine. For behavioral experiments, there was a minimum of four weeks between viral surgery and baseline short-access cocaine self-administration to allow for sufficient removal of KOR from transfected neurons.

#### Catheter Implantation

Indwelling catheters were assembled in-house using custom 26G guide cannulas with a 5mm up/bent projection on one side and a 6mm projection cut below the pedestal purchased from Plastics One (8IC315GFLX03), Liveo laboratory tubing (cat 508-001), and silicone. Catheters were cut at different lengths for males (10cm) and females (8cm) to accommodate for differences in size.

If multiple surgeries, the catheter implantation surgery was performed second to reduce amount of time necessary to maintain patency. The catheter tubing was inserted into the right jugular vein, secured with sutures, and the catheter pedestal was secured onto the dorsal side for back-mount access. At the conclusion of surgery, the catheter was back-filled with a 60% polyvinylpyrrolidone-40 solution in saline containing gentamicin (20mg/mL) and heparin (1,000units/mL) and capped with an in-house fabricated dust-cap (PE20 tubing). Rats were allowed to recover for 7-9d before starting flushing their catheters daily with approximately 0.05mL of 0.9% saline or heparinized saline (80units/mL) and beginning cocaine self-administration.

### Drugs and Viral Vectors

Norbinaltorphimine (norBNI) was suspended in ACSF (5μg/0.3μL) and injected bilaterally into the NAc (0.3μL/hemisphere) two days prior to the first long-access SA session. U69,593 (Sigma Aldrich) was dissolved in solvent 0.01% DMSO and aCSF for a final concentration of 1μM. Cocaine hydrochloride was dissolved in physiological saline (5mg/mL), filtered, and administered at a dose of 5mg/kg.

The following viral constructs were manufactured by the University of Washington Center for Excellence in Neurobiology of Addiction, Pain, and Emotion’s Molecular Genetics Resource Core: AAV1-FLEx-ChR2-eYFP, AAV1-FLEx-SaCas9-U6-HA-sgOprk1, AAV1-FLEx-SaCas9-U6-HA-sgGabrg2 Control, and AAV1-Cre-GFP. Titer for all viruses range from 0.5 to 5x10^9^ particles.

### Cocaine Self-Administration

All behavioral experiments were performed in Med Associates modular operant chambers outfitted with a liquid swivel attached to a syringe pump, two nosepoke ports with internal cue lights, a houselight, a tone generator, and a white noise generator. The houselight and white noise turned on to mark the beginning of the session and availability of cocaine. Operant responding into the assigned active nosepoke port resulted in one cocaine infusion (FR1, 5mg/mL/kg at 0.02755mL/sec) and the start of a 20-second time-out period during which the houselight and white noise are off, an audiovisual cue is on (tone + nosepoke port cue light), and further operant responses are recorded but do not result in additional cocaine infusions. The second nosepoke port (referred to as inactive nosepoke) does not have any programmed consequences and is used to assess general locomotion and responding behavior.

Rats are first trained on short-access self-administration (SA) sessions that last for one hour per session until they reach our learning criterion (ten active responses per session for three consecutive sessions). After reaching criterion, rats continue short-access SA for five additional days to establish a baseline of cocaine consumption. Then, rats enter two five-day blocks of long-access (six-hour sessions) SA. Between each block (short-access block, 1^st^ long-access block, and 2^nd^ long-access block), the rats receive a 0- to 2-day break from SA. At the conclusion of behavioral experiments, we performed a linear regression on active responses during the first hour of each session across sessions and classified rats with a significant, positive correlation as escalators and rats with a non-significant or significant, negative correlation as non-escalators, as previously described^20^.

### Microinjections

We injected norBNI (5μg/mL) or vehicle bilaterally into the NAcC two days prior to the rats’ first long-access SA session and at least one day post their last short-access SA session. We inserted a bilateral injector with 1mm projection (Plastics One, part#: 8IC235ISPCXC) into the guide cannula and used two Hamilton syringes (1801RN 10ul SYR (26s/2”/2), ref#: 84877) coupled with a kdScientific injector (model 210) to inject 0.3μL/hemisphere followed by a two-minute wait period before removing the injector.

### Ex Vivo Fast-Scan Cyclic Voltammetry

For slice voltammetry experiments animals were euthanized with pentobarbital (150mg/kg) and perfused with a cold sucrose solution. The brain was removed and 300μm coronal slices containing the NAcC were prepared in an oxygenated sucrose solution. Slices were held in a recovery chamber containing oxygenated aCSF for at least 45min at 32°C before any recordings took place. To record dopamine transients the slices were placed in a recording chamber, and continually perfused with oxygenated aCSF at 33°C. Carbon-fiber microelectrodes (CFM) were fabricated with a fused-silica capillary^21^ and cut to have a recording surface of ∼175μm. Transients were evoked by a single pulse of a blue LED through the microscope objective. Recordings were accomplished by applying a triangular waveform in which, the CFM potential is ramped from -0.4V (versus Ag/AgCl) to +1.3V and then back -0.4V at a rate of 10 Hz. The experiment did not begin until dopamine recordings were stable for 10min. Evoked transients during baseline and experimental recordings took place every two minutes. Following stable recordings an additional 5 recordings were taken for baseline, then 1μM of U69,593 within oxygenated ACSF was washed on and an additional ten recordings were performed. Waveform generation, data acquisition, and analysis were carried out using two PCI cards and software written in LabVIEW 7.1 (National Instruments, Austin, TX).

### Histology

For all behavioral experiments, rats were euthanized with pentobarbital (150mg/kg) and transcardially perfused with saline followed by 4% paraformaldehyde. For cannulated rats, 0.3μL of blue dye was injected into the NAcC prior to perfusion. Their brains were removed and submerged in ascending concentrations of sucrose, ending with 30% sucrose in PBS, before being frozen and sliced into 40μm coronal sections on a cryostat. Slices from cannulated rats were mounted and the location of the blue dye was imaged and analyzed to confirm location of pharmacological manipulations. Slices from virally-injected animals underwent immunohistochemistry for HA-tag to confirm location of viral transduction of the CRISPR virus. Rats with placements outside of the tar were excluded.

### Statistical Analysis and Data Visualization

Statistical analyses and data visualization for all behavioral experiments were performed in R (version 4.0.5). Mixed effects models were computed with the “lme4” package (version 1.1.27.1). ANOVA was computed with the “afex” package (version 1.0.1). HSD was computed with the “emmeans” package (version 1.7.0). Linear regression was computed with base R. Data visualization was computed with “ggplot2” (version 3.3.5). Figure compilation was performed in Adobe Illustrator (CS4).

## RESULTS

### Pharmacological Inhibition of KOR in the NAcC Blocks Cocaine Escalation

To test the necessity of KOR activation in the NAcC for escalation of cocaine consumption, we pharmacologically inhibited KOR via local microinjection of the long-lasting KOR antagonist, norBNI, prior to transitioning rats from short-access SA to long-access SA (Fig. 1b). A mixed effects model analysis revealed an interaction between treatment (vehicle vs norBNI) and time (SA sessions) demonstrating differential cocaine consumption patterns over SA across the two groups (Fig. 1c). As a group, vehicle-treated rats escalated in their cocaine consumption over long-access SA sessions, as expected, while norBNI-treated rats did not escalate in their consumption (Fig. 1d). Analyzing average cocaine consumption per block, vehicle-treated rats show increased responding in the first and second block of long-access SA (LgAb1 and LgAb2, respectively) in comparison to their short-access baseline (ShA) while norBNI-treated rats showed no significant difference in responding in LgAb1 or LgAb2 compared to ShA (Fig. 1e). However, norBNI-treated rats did show a significant increase in responding from LgAb1 to LgAb2, potentially indicative of a reduction in the effectiveness of the norBNI injection over the two weeks or a secondary mechanism behind escalation emerging. Categorizing rats as escalators or non-escalators based on a linear regression analysis of drug consumption across SA session where significant, positive correlations are deemed escalators, demonstrated a trend toward a lower proportion of escalators in the norBNI-treated group (Fig. 1f). Overall, pharmacologically inhibiting KOR in the NAc prior to long-access SA blocked (or at least prolonged) the development of the SUD-phenotype of escalation.

**Fig 1.**
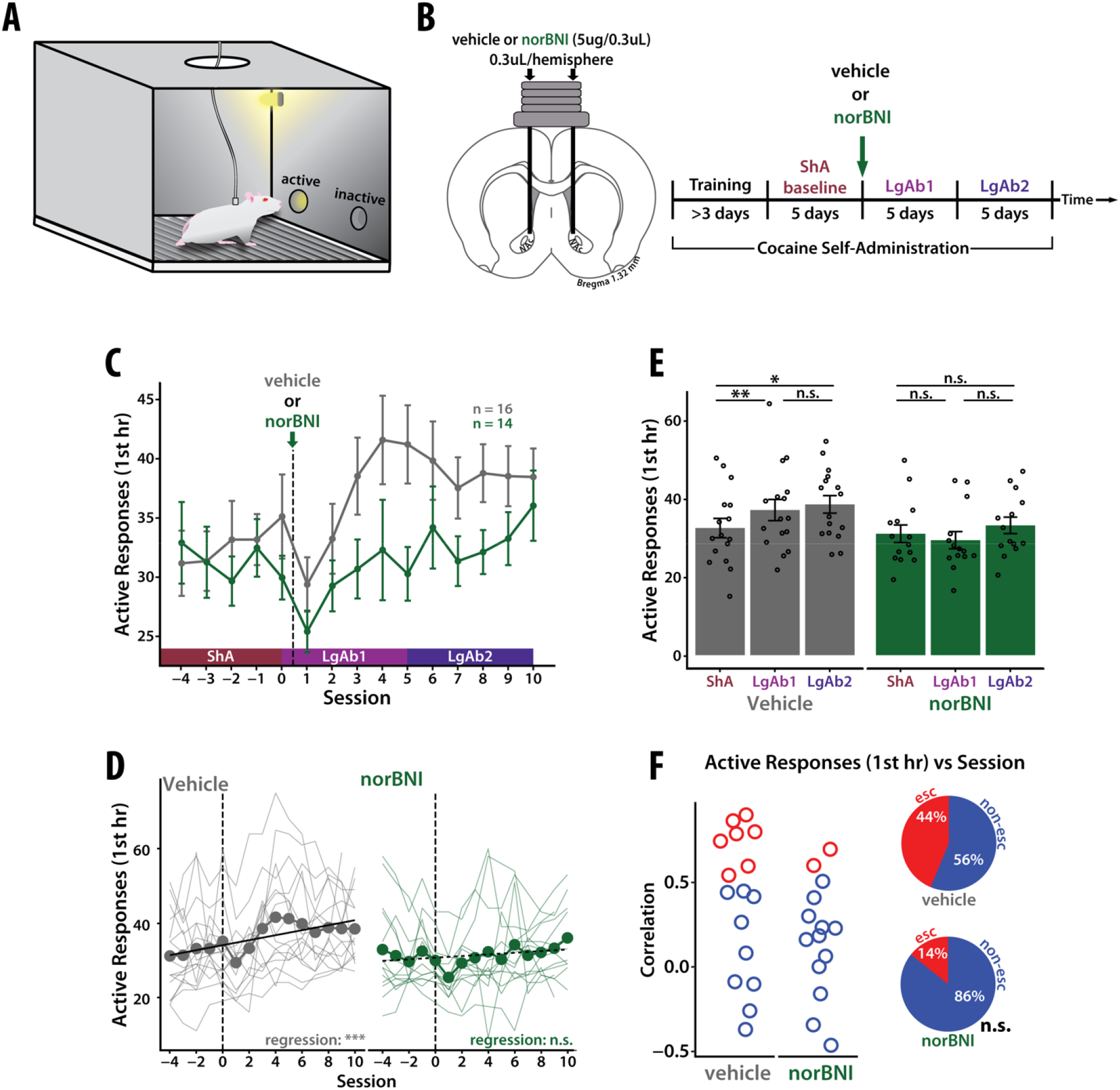
Pharmacological Inhibition of KOR in the NAcC Blocks Cocaine Escalation. **(A)** Cocaine self-administration setup. **(B)** experimental design. **(C)** Local NAcC infusion of long-lasting KOR antagonist, norBNI, prior to long-access SA results in differential drug-taking across sessions compared to control (main effect of session F(14,368)=4.78, p-value < 0.001; main effect of group F(1,28)=2.53, p-value > 0.05; interaction F(14,368)=2.00, p-value < 0.05). Active responses for only the first hour of SA and error bars show SEM. **(D)** Thinner lines represent individual rats. Black line indicated line of best fit from a linear regression. As a group, vehicle-treated rats showed an increase in active responses across session (r^2^=0.058, p-value < 0.001) whereas norBNI-treated rats did not (r^2^=0.009, p-value > 0.05). **(E)** Active responses during the first hour of SA averaged into three 5-session blocks: short-access (ShA), long-access block 1 (LgAb1), and long-access block 2 (LgAb2). A two-way anova revealed a significant interaction between group and block (F(2,56)=4.49, p-value < 0.05). Post-hoc Tukey test showed a significant increase between ShA and LgAb1 for vehicle-treated rats (p-value < 0.01) and between LgAb1 and LgAb2 for vehicle-treated rats (p-value < 0.05). No significant changes were detected in norBNI-treated rats. **(F)** Each rat was classified as an escalator or a non-escalator based on a linear regression analysis of active responses across session. Rats with a significant, positive regression are classified as escalators and rats with a non-significant or a significant, negative regression are classified as non-escalators. (left) correlation values of escalators and non-escalators in the two groups. (right) percentage of escalators to non-escalators in both groups. A Chi-squared goodness of fit test showed no significant difference between the two groups (p-value > 0.05).

### Virally-mediated CRISPR Knock-Out of oprk1 in the VTA Eliminates KOR-mediated Inhibition of Dopamine in the NAcC

To knock-out KOR on specific neurons/terminals in the NAcC, we developed a viral CRISPR-SaCas9 construct targeting the *oprk1* gene (encoding for KOR) to knock-out KOR in specific neuronal populations (Fig. 2a). To confirm the knock-out of functional KOR in neurons, we injected the viral CRISPR-SaCas9 construct into the VTA and, after incubation, measured the effect of KOR agonist U69,593 on stimulated dopamine release in the NAcC with *ex vivo* fast-scan cyclic voltammetry (FSCV). We additionally injected channelrhodopsin into the VTA to enable light-activation of VTA projections in the NAcC with simultaneous recording. Bath application of U69,593 had a differential effect on evoked dopamine release across groups (Fig. 2c). An unpaired t-test revealed a significant decrease in the effect of U69,593 on evoked dopamine release in VTA-oprk1-KO rats compared to control rats (Fig. 2c). Our virally-mediated CRISPR knock-out of *oprk1* was sufficient to significantly reduce the ability of a KOR agonist to modulate dopamine release in the NAcC when expressed in the VTA demonstrating the functional validity of our construct.

**Fig 2.**
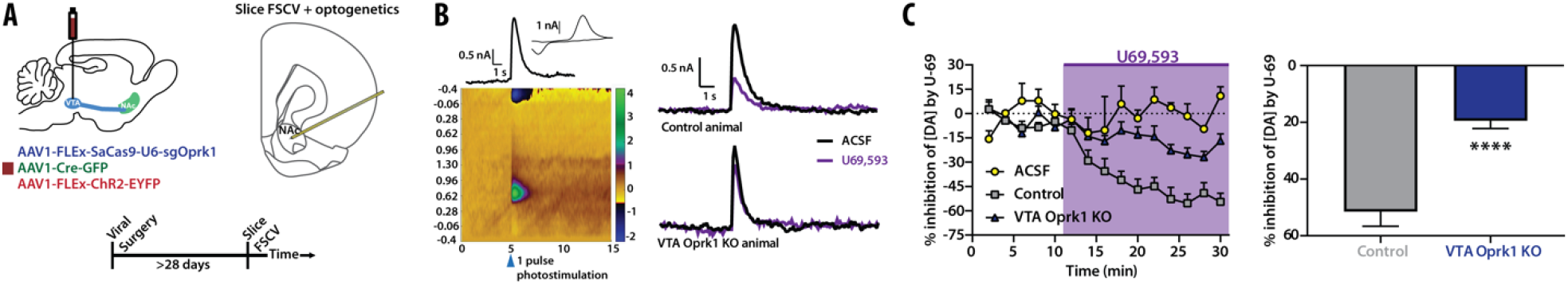
Virally-mediated CRISPR-SaCas9 Targeting *oprk1* Effectively Knocks Down KOR Function. **(A)** Experimental setup. **(B)** (left) Expression of ChR2 was sufficient in stimulating dopamine transients following a pulse of blue light onto a coronal slice. current vs time (top left), the corresponding CV (top right), and a color plot with current (color), time (x-axis), and potential (y-axis). (right) Representative traces of stimulated dopamine release before and after washing the KOR agonist U69,593, in both a control animal (top) and an animal with SaCas9 mediated KOR knock-out (bottom). **(C)** (right) Time series of stimulated dopamine release under various conditions. A subset of slices underwent 15 stimulations without the U69,593 being washed on (yellow circles, n=3). The other two groups, control (gray squares n=7) or KOR knock-outs (purple triangles n=9), had 5 stimulations (10 mins), before U69,593 was applied to the ACSF bath (purple box). A mixed-effects model revealed there was a main effect of group (F(2,16)=12.81, p-value <0.001), and there was an interaction of group by time (F(28,216) = 6.299, p<0.0001). A post-hoc Tukey test showed there was a significant difference between the KOR knock-outs and control (p-value <0.0001), and a difference between the KOR knock-outs and ACSF (p-value <0.0001). (left) To test the normalized change in dopamine release after U69,593 application, the 5 stimulations just prior to U69,593 wash on were used as a baseline, the last 5 stimulations before experimental end were averaged and used to identify the % inhibition from baseline. We observed a significant differences in the effect a KOR agonist had on inhibition of dopamine release (unpaired t-test (t(14) = 5.87, p<0.0001).

### Virally-mediated CRISPR Knock-Out of oprk1 in the VTA but not the NAcC Blocks Cocaine Escalation

To delineate which KOR-expressing neurons in the NAcC are necessary for cocaine escalation, we knocked-out (KO) KOR in NAc neurons or VTA neurons independently using virally-mediated CRISPR-SaCas9 technology targeting the *oprk1* gene prior to training the rats on the cocaine SA. A control CRISPR virus was injected either into the NAcC or VTA (Fig. 3a). There was no difference in cocaine consumption between the two groups of control-virus rats (supp fig. 1), and therefore, the two groups were pooled. All the rats, regardless of treatment, learned cocaine SA and there was no significant difference in baseline cocaine consumption during short-access SA (supp fig. 2). A mixed effects model analysis revealed a significant interaction between group (virus/location) and time (SA sessions) demonstrating differential cocaine consumption patterns over SA across the groups (Fig. 3c). As a group, control-virus rats and NAcC-oprk1-KO rats escalated in their cocaine consumption over long-access SA sessions, while VTA-oprk1-KO rats did not escalate in their consumption (Fig. 3d). Analyzing average cocaine consumption per block (Fig. 3e), control-virus rats showed increased responding from ShA to LgAb2; NAcC-oprk1-KO rats showed increased responding from ShA to LgAb1; VTA-oprk1-KO rats did not show any significant differences across all three blocks. Examining the proportion of escalators vs non-escalators across groups demonstrated a significant reduction in the proportion of escalators in VTA-oprk1-KO rats compared to control-virus rats, with no significant difference in the proportions of escalators in NAcC-oprk1-KO rats compared to control-virus rats (Fig. 3f). Additionally, there was no interaction between session and sex for any group, although control-virus and VTA-oprk1-KO groups did have a main effect of sex (supp fig. 3). Overall, the knock-out of *oprk1* from VTA neurons (including their presynaptic terminal in the NAcC), but not from medium spiny neurons in the NAcC, was sufficient to completely block escalation of cocaine consumption. This demonstrates that activation of KOR on VTA neurons is necessary specifically in the transition from casual to SUD-like cocaine consumption.

**Fig 3.**
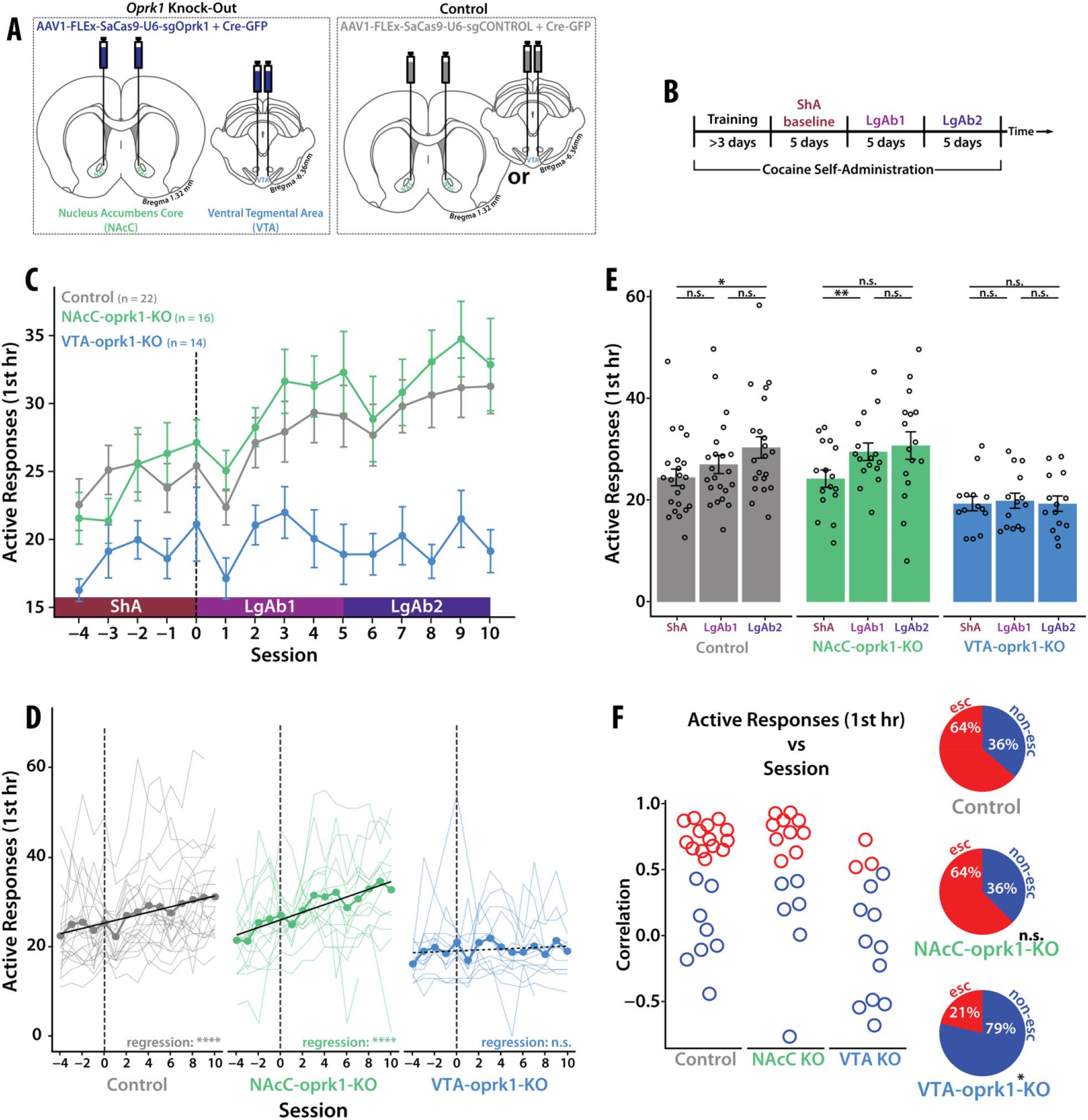
CRISPR-SaCas9 Knock-Out of KOR in VTA Neurons but not in NAcC Neurons Blocks Cocaine Escalation. **(A)** Viral strategy. **(B)** Behavioral timeline. **(C)** A mixed effects model on active responses in the first hour across session revealed a significant main effect of session (F(14,657)=10.60, p-value < 0.001), main effect of group (F(2,49)=7.47, p-value < 0.01), and an interaction (F(28,657)=2.10, p-value < 0.001). **(D)** Thinner lines represent individual rats. Black line indicated line of best fit from a linear regression. As a group, control-virus rats and NAcC-oprk1-KO rats showed an increase in active responses across session (control-virus: r^2^=0.07, p-value < 0.001; NAcC-oprk1-KO: r^2^=0.14, p-value < 0.001;) whereas VTA-oprk1-KO rats did not (r^2^=0.004, p-value > 0.05). **(E)** Active responses during the first hour of SA averaged into three 5-session blocks: short-access (ShA), long-access block 1 (LgAb1), and long-access block 2 (LgAb2). A two-way anova revealed a significant interaction between group and block (F(4,98)=2.97, p-value < 0.05). Post-hoc Tukey analysis demonstrated the following effects: (control-virus) a significant increase between ShA and LgAb2 (p-value < 0.05); (NAcC-oprk1-KO) a significant increase between ShA and LgAb1 (p-value < 0.01); (VTA-oprk1-KO) no significant differences. **(F)** Each rat was classified as an escalator or a non-escalator based on a linear regression analysis of active responses across session. Rats with a significant, positive regression are classified as escalators and rats with a non-significant or a significant, negative regression are classified as non-escalators. (left) correlation values of escalators and non-escalators in the two groups. (right) percentage of escalators to non-escalators in both groups. A Chi-squared goodness of fit test showed a significant difference between the control-virus and VTA-oprk1-KO (p-value < 0.05) but not NAcC-oprk1-KO (p-value > 0.05).

## DISCUSSION

Here, we demonstrated that local inhibition of KOR via a long-lasting antagonist, norBNI, in the NAcC blocks cocaine escalation in a long-access model of cocaine SA. To probe which KOR in the NAcC are responsible for this effect, we validated a novel virally-mediated CRISPR targeting the *oprk1* gene and utilized this technique to knock-out KOR of NAcC medium spiny neurons and VTA neurons. We demonstrated that knocking-out KOR in VTA neurons, but not NAcC neurons, completely blocked cocaine escalation. We did not observe effects on learning or baseline cocaine consumption when KOR was knocked out of VTA neurons indicating that this effect is specific to escalation behavior. Therefore, the development of escalation (a SUD-like phenotype), but not cocaine consumption in general, requires activation of KOR on VTA terminals in the NAcC.

It is important to note that KORs are also expressed on VTA cell bodies and presumably VTA projections to regions other than the NAcC^17,22^. The viral technique used in this paper was not specific to the VTA-NAcC pathway and there is yet a well-validated technique to knock-out receptors selectively on terminals. Therefore, it is possible that our effect could be due to knocking out KOR in areas other than VTA terminals in the NAcC. However, in combination with the pharmacological data, which only affects KORs present in the NAcC, the data is highly suggestive that it is KOR on VTA pre-synaptic terminals in the NAcC necessary our observed effect on escalation of cocaine consumption.

The projection from the VTA to the NAcC mostly consists of dopamine neurons^23^. As demonstrated here and in previous literature^13^, activation of KOR inhibits dopamine release. Knocking out KOR from VTA neurons blocks dynorphin activation of KOR in the NAcC and the resulting decrement in dopamine transmission. Previous work has demonstrated that decreases in response-contingent cue-evoked dopamine release across long-access cocaine SA is causal to escalation of cocaine consumption^20^. Therefore, it is possible that in these experiments, blocking the activation of KOR on VTA neurons through pharmacology or gene-editing inhibited KOR from decreasing dopamine responses ultimately resulting in a lack of escalation.

The results of this paper provide evidence for KOR activation on VTA terminals in the NAcC to drive escalation of cocaine consumption, potentially working upstream of dopamine through KOR-activated inhibition of dopamine release. KOR has been implicated in SUD of various addictive substances including cocaine^2,24–28^, methamphetamine^19^, heroin^18^, and alcohol^6,29,30^. Therefore, these results will likely be relevant to more than cocaine use disorder alone. Uncovering the neural mechanisms behind SUD will inform therapeutic strategies to combat the current drug epidemics.

## ACKNOWLEDGEMENTS

We thank Jacqueline McAleer for technical assistance. This work was funded by the National Institutes of Health grants F31-DA048562 (RF), T32-DA007278 (PP), R01-DA039687 (PP), and R37-DA051686 (PP).

## SUPPLEMENTAL

**Supp Fig 1.**
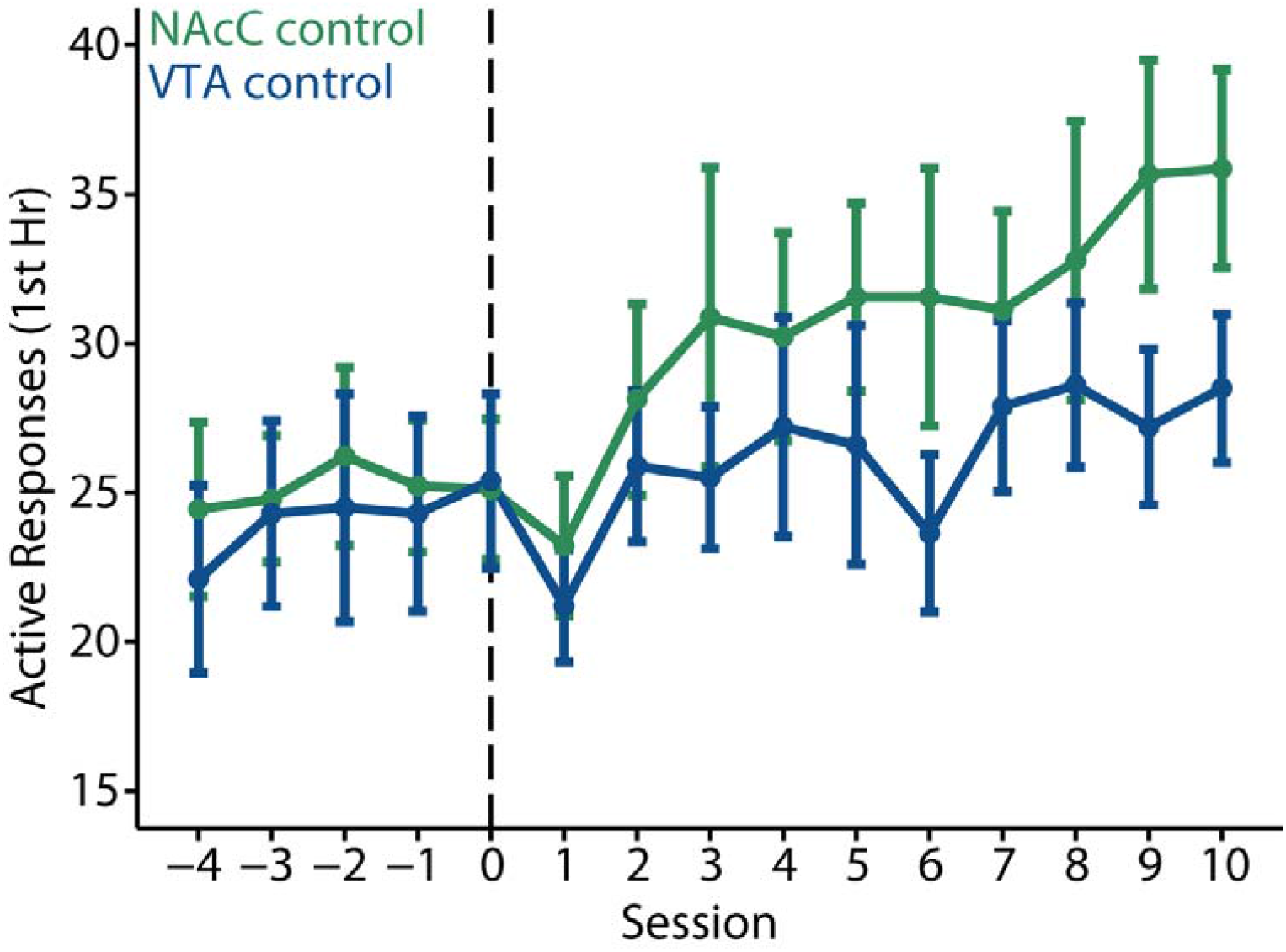
Location of CRISPR Control Virus Injection Does Not Impact Cocaine SA. Active responses across session for all control-virus animals, divided by their injection region. A mixed effect model analysis revealed no main effect of group nor interaction.

**Supp Fig 2.**
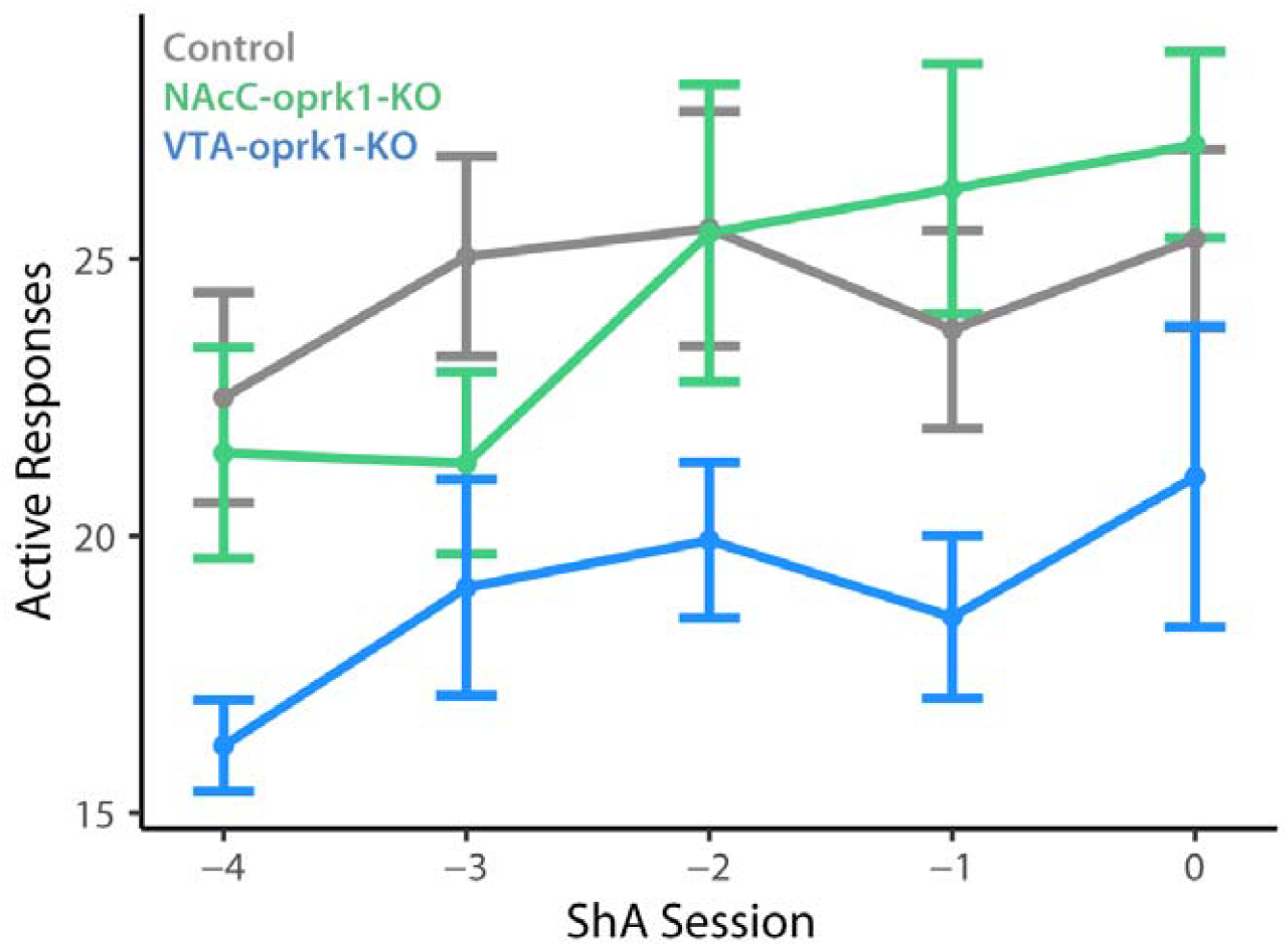
CRISPR KO or KOR does not Affect Baseline Cocaine Consumption. Active responses during 5 baseline short-access (ShA) cocaine SA sessions. A mixed effects model showed no significant main effect of group (p-value > 0.05) nor interaction (p-value > 0.05).

**Supp Fig 3.**
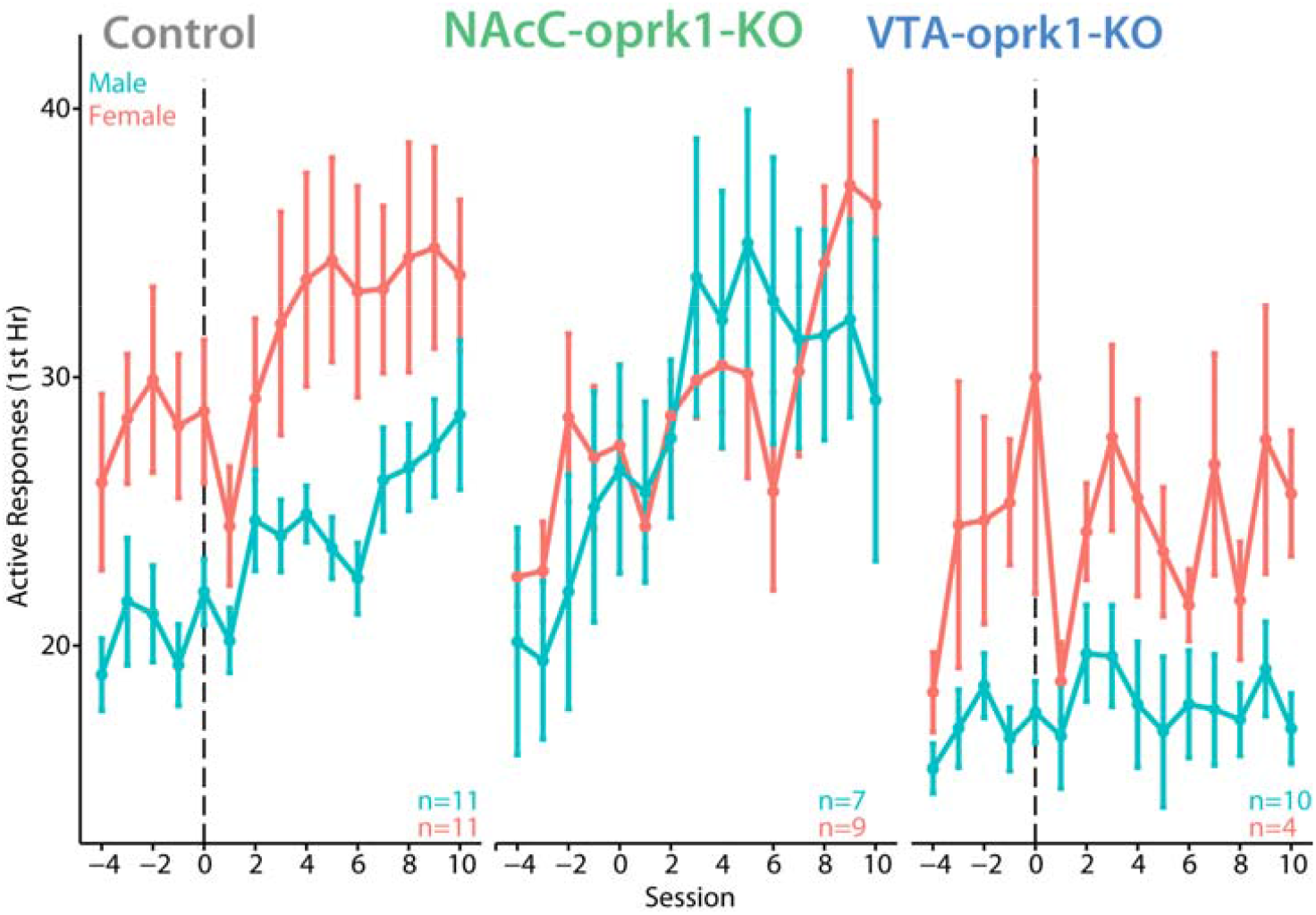
CRISPR KO of KOR Effect on Cocaine SA in Males and Females. For control-virus rats, a mixed effects model revealed a main effect of session (F(14,275)=7.67, p-value < 0.001) and a main effect of sex (F(1,20)=6.20, p-value < 0.05), but no interaction. For NAcC-oprk1-KO rats, a mixed effects model revealed only a main effect of session (F(14,183)=6.70, p-value < 0.001). For VTA-oprk1-KO rats, a mixed effects model revealed only a main effect of sex (F(1,12)=10.4, p-value < 0.01).

